# VisVariant: A java program to visualise genetic variants in next-generation sequencing data

**DOI:** 10.1101/2021.02.12.431037

**Authors:** King Wai Lau, Michelle Kleeman, Caroline Reuter, Attila Lorincz

## Abstract

**Summary:** Extremely large datasets are impossible or very difficult for humans to comprehend by standard mental approaches. Intuitive visualization of genetic variants in genomic sequencing data could help in the review and confirmation process of variants called by automated variant calling programs. To help facilitate interpretation of genetic variant next-generation sequencing (NGS) data we developed VisVariant, a customizable visualization tool that creates a figure showing the overlapping sequence information of thousands of individual reads including the variant and flanking regions.

**Availability and implementation:** Detailed information on how to download, install and run VisVariant together with an example is available on our github website [https://github.com/hugging-biorxiv/visvariant].

## Introduction

Next-generation sequencing (NGS) is fast becoming the de facto tool to identify genetic variants. Many variants can be found in one sequencing experiment. The true positive rate of automatically-called variants depends on multiple factors including read depth, flanking sequence, systematic bias of sequencing platform and variant calling algorithms. Visualization of automatically called variants with flanking regions could improve the confidence of the calls. Another way to increase the confidence is by superposing the NGS reads generated from multiple samples with the same phenotype.

Multiple existing NGS alignment visualization tools can be used to visualize genomic regions around variants. These tools are capable of displaying aligned reads with one or more samples at the same time, such as IGV (Robinson et al., 2011), svviz (Spies et al., 2015), tview (Li et al., 2009), BamView (Carver et al., 2013), Tablet (Milne et al., 2010), gbrowse2 (Stein et al., 2013), Savant (Fiume et al., 2012) and ZOOM Lite (Zhang et al., 2010).

However, there is a trade-off between visualising the fine details of read sequences and zooming out to visualize many reads together and these tools do not perform very well when the read depth of a sample increases or when visualising multiple samples at once. In addition, these tools will find it difficult to visualisation of more than, say 10 samples at the same time and do not allow grouping the multiple samples together. To circumvent these issues, we developed a customizable visualization tool, VisVariant, to help facilitate interpretation of genetic variants from NGS data.

## Implementation

### Main features

VisVariant is an adaptation of ALVIS (Roland et al., 2016), a visualization tool for multiple protein sequence alignment. Our adaptation is able to maintain the read sequence information while summarizing many reads together. This method transforms a read sequence into an undirected path, where the nucleotide is a node and adjacent nucleotides are connected by an edge. The resulting paths covering the variant are summarized into a two-dimensional graph where each row represents the nucleotide of the reads and the columns represent a segment of the reference genome, e.g. the genomic coordinate of the variant +/− flanking region. This two-dimensional graph is displayed on a fixed size panel (Figure 1).

**Fig. 1:**
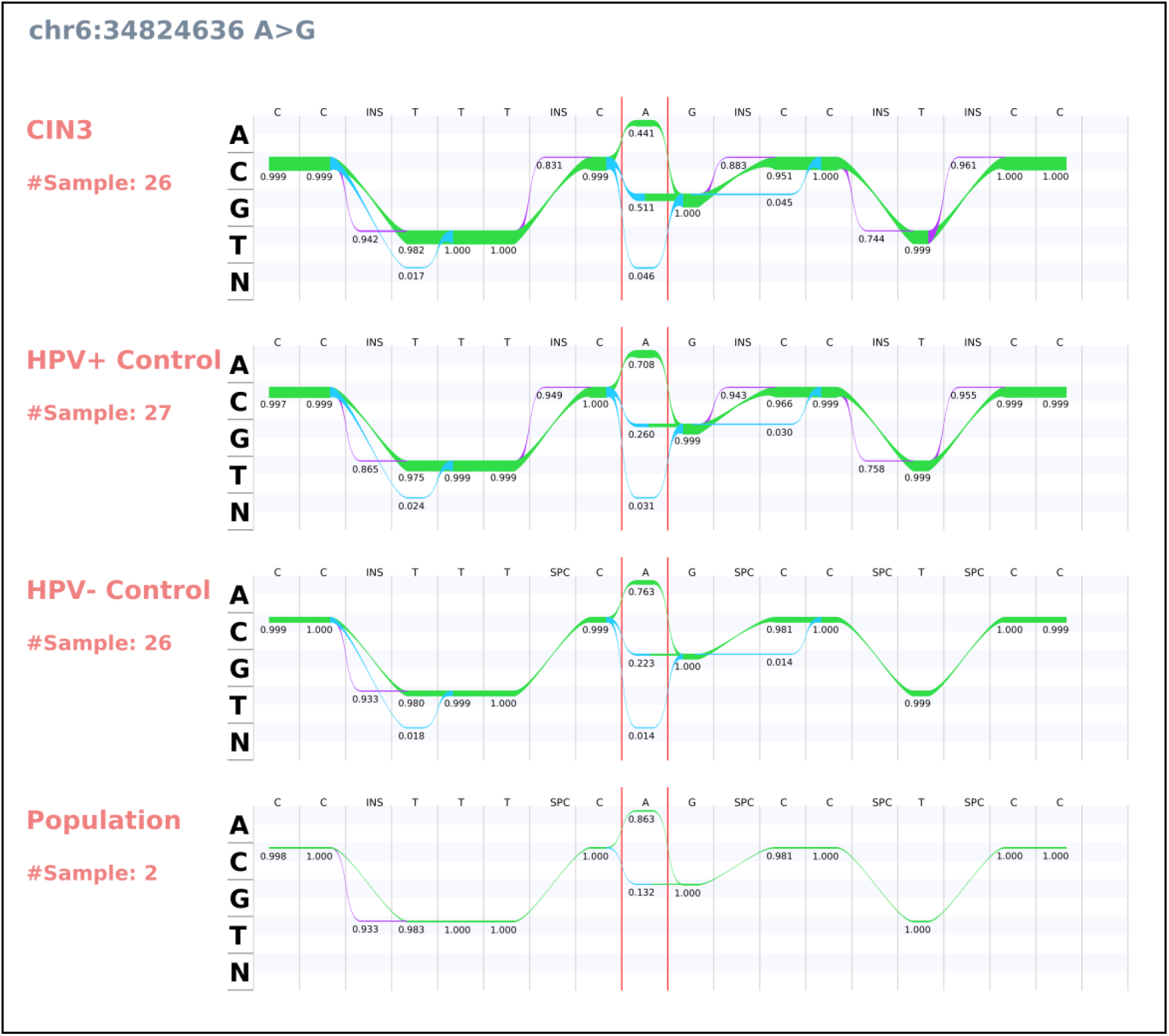
A typical VisVariant figure. This example shows a variant rs11755393 on chromosome 6 at position 34824636 and its flanking regions. There are four panels, one for each phenotype, CIN3, HPV+ control, HPV-control and population. The variant is delimited by two continuous vertical red lines. The number in the cells indicate the variant allele frequency. The thickness of the trace is proportional to the total number of reads covering that nucleotide. When the sequenced nucleotide matches the reference, the trace colour is green. Insertion are represented in purple. Any other changes from the reference are represented in blue. In order to align the variant across the panels, one or more space columns (SPC) may be inserted. Note that some of the edges are filtered out with (readcountrenderthreshold=25) to reduce the number of edges in this figure.

The advantage of our approach is: 1) Allowing visualization of the details of the read sequence information via the two dimensional graph; 2) Displaying the summary statistics of any number of reads on the graph; 3) Displaying multiple groups of samples together using multiple panels; 4) Identifying difficult variant calling regions, such as repeated regions and homopolymer regions, because they tend to increase the number of edges on the two dimensional graph; 5) Identifying systematic biases, such as a variant induced by the sequencing platform can be found by comparing multiple panels altogether.

VisVariant requires four user input files: 1) A VCF file for a list of called variants; 2) A genomic reference FASTA file where NGS reads are aligned; 3) A phenotype file describing the BAM file name of each sample and group name; and 4) A parameter file to set up the parameters of VisVariant. In addition, VisVariant requires two directory paths, the first for the location of a set of BAM files and the second for saving the resulting figures. A VisVariant run takes approximately 20 seconds on a 3.4 GHz Intel(R) Core(TM) i7 processor and 8 GB RAM to create a figure for a variant with 81 samples at median read depth 150x per sample. The resolution of the resulting figure is of high quality and can be used in journal publication, but clarity also depends on the computer screen resolution.

Figure 1 shows an example of a variant across multiple samples in multiple groups with one panel for each group. The panels are organized by the order of the group name in the phenotype file. Within the panel, a matrix structure is shown, where the rows correspond to the nucleotides of the read and the columns correspond to the nucleotides of the reference. An extra row at the bottom (N) of the panel is used when a nucleotide is deleted from the reads compared to the reference. A column (INS) is added when a nucleotide is inserted to the read. One or more space (SPC) column(s) may be added such that the panels are aligned to each other. The variant is delimited by the red vertical lines. The number in the cell shows the allele frequency of the corresponding nucleotide.

The figure settings can be specified in the VisVariant command line or in a parameter file. The user can change the default colours (readcolour_unvaried, readcolour_varied and readcolour_insertion), the width and height of the cells (columnwidth and rowheight, respectively), maximum number of columns (columns), number of base pairs for the flanking regions (flank) and filtering of edges when one node or both nodes do not have enough read depth (readcountrenderthreshold). In addition, the user can change the width (width) and height (height) to control the resolution.

### Installation and usage

VisVariant was written in Java and uses the samtools HTSJDK library (http://samtools.github.io/htsjdk/) for reading BAM files and a VCF file. Installation of VisVariant is straightforward. Download the compressed file from our github repository [https://github.com/mel-bioinfo/VisVariant] and unzipping the file into a folder, the user will find a jar file for the VisVariant program and an example parameter file. Users can also download an example dataset extracted from Genome in a Bottle. VisVariant supports command line options and a parameters file with one line per option. A readme file explains the options and the exact command is documented in the README.md file.

### Functionality

VisVariant functionality is best illustrated using in-house generated NGS data by the Ion Proton sequencing platform (Figure 1). In this dataset, we have sequenced the whole exome of 81 human samples. The samples can be divided into four groups: 1) Population (n = 2); 2) HPV-Negative Control (n = 26); 3) HPV-Positive Control (n = 27); 4) Cervical intraepithelial neoplasia grade 3 (CIN3) (n = 26) the disease outcome of interest. The HPV-Positive Control group comprised base-line samples for women who had an HPV infection but who did not progress to CIN3. The HPV-Negative Control group included base-line samples for women who did not have an HPV infection and did not progress to CIN3. The Population group comprised cell lines purchased from the Coriell Institute (NA12878).

Figure 1 shows the visualization of an A to G variant rs11755393, located on chromosome 6 at position 34824636. In this figure, there are four panels, one for each disease group. The order of the panel reflects the progression of disease from no disease at the bottom to CIN3 at the top. The allele frequency of the alternative allele (G) increases from 0.132 (no disease) to 0.511 (CIN3), matching the expectation of disease progression. We can be confident that this variant is true positive because there is no homopolymer sequence in the flanking region immediately adjacent to the variant and the figure does not have many edges indicating this region is not difficult to sequence.

We used VisVariant to compare the sequences of the European population cell line NA12878 on four sequencing platforms (Supplementary Figure 1), IonProton, Illumina, Complete Genomics and PacBio. The BAM files for Illumina (n=1), Complete Genomics (n=2) and PacBio (n=1) were downloaded from the Genome in a Bottle (Zook et al., 2014) FTP site (ftp://ftp-trace.ncbi.nlm.nih.gov/giab/ftp/data/NA12878/). The BAM file for the Ion Proton were obtained the Population group above. Looking at position rs11755393 again (Supplementary Figure 1), we showed that all platforms performed equally well and called an A at that position for NA12878. However, PacBio and IonProton produced an insertion and/or deletion five bases upstream of rs11755393, an error probably due to the homopolymer sequence in the region. Test data can be found in [https://github.com/hugging-biorxiv/visvariantData].

## Conclusions

We present an easy-to-use visualization tool that can intuitively display specific nucleotide variants in NGS data. This allows human operators to visually review and to confirm the most convincing variants from others that may be related to sequencing errors or errors in the validation process. We expect our tool will find many uses in the burgeoning genomics field allowing better visualization of nucleotide variants of specific interest.

