## Supplemental Figure 1 for "VisVariant: A java program to visualise genetic variants in next-generation sequencing data"

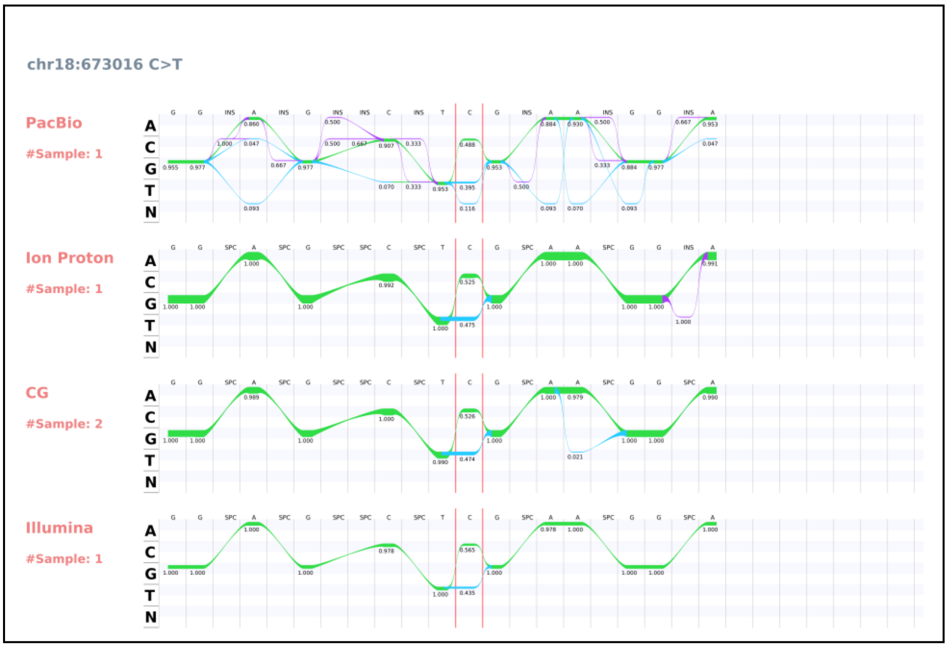


**Supplementary Figure 1.** VisVariant shows a variant rs11755393 and its flanking regions sequenced using four different sequencing platforms, PacBio, Ion Proton, Complete Genomics and Illumina.
